# Functional connectivity between interoceptive brain regions is associated with distinct health-related domains - a population-based neuroimaging study

**DOI:** 10.1101/2022.07.27.500935

**Authors:** A Luettich, C Sievers, F Alfaro Almagro, M Allen, S Jbabdi, SM Smith, KTS Pattinson

## Abstract

Interoception is the sensation, perception, and control of signals from within the body. It has been associated with a broad range of physiological and psychological processes. Further, interoceptive variables are related to specific regions and networks in the human brain. However, it is not clear whether or how these networks relate empirically to different domains of physiological and psychological health at the population level.

We analysed a dataset of 19 020 individuals (10 055 females, 8 965 males; mean age: 63 years, age range: 45 – 81 years), who have participated in the UK Biobank Study, a very large scale prospective epidemiological health study. Using canonical correlation analysis (CCA), allowing for the examination of associations between two sets of variables, we related the functional connectome of brain regions implicated in interoception to a selection of non-imaging health and lifestyle related phenotypes, exploring their relationship within modes of population co-variation.

In one integrated and data driven analysis, we obtained four statistically significant modes. Modes could be categorised into domains of affect and cardiovascular health, breathing, obesity, and subjective health (all p < 0.0001) and were meaningfully associated with distinct neural circuits.

Circuits represent specific neural “fingerprints” of functional domains and set the scope for future studies on the neurobiology of interoceptive involvement in different lifestyle and health related phenotypes. Therefore, our research contributes to the conceptualisation of interoception and may lead to a better understanding of co-morbid conditions in the light of shared interoceptive structures.

## Introduction

Interoception is defined as the sensing, interpretation, integration and regulation of internal states for the maintenance of homeostasis ^1–4^. Regulation of cardiac, digestive and respiratory function are closely related to interoceptive processes ^2^, as are emotional ^5–10^, cognitive ^11–15^ and reflexive, self-related states ^16–18^. Thus, interoception links “the mind” with “the body”.

Interoceptive ability is potentially a key mediator of health-relevant variables, such as individual differences in physiological functioning and symptom perception. Disturbed interoception is implicated in somatic, developmental, neurological, neurodegenerative, and notably also psychiatric conditions including for instance disorders of self-awareness, anxiety and depression ^19–21^.

Currently, it is far from clear whether or how specific neural networks relevant to interoception relate empirically to different physiological, psychological and generally health-related domains, i.e., whether they show sensitive individual “fingerprints”, at the population level. To date no large-scale studies have been done on this. Thus, we utilised a large scale database, the UK Biobank Imaging Study ^22^, and related the functional connectome of a selection of brain regions relevant to interoception to a selection of lifestyle and health related phenotypes in order to explore their relationship within modes of population co-variation ^23^.

## Methods

### Participants

Imaging and non-imaging data from the UK Biobank cohort was available from a total of 19,020 participants (10,055 females, 8,965 males). The average age of participants was 63 years (± 7 standard deviations, range: 45 - 81).

### Imaging

#### Acquisition

Low sample size is a common problem in neuroscience research, impacting reproducibility and scientific progress ^24^. UK Biobank is a population level prospective epidemiological health study collecting extensive behavioural, physiological and general health related information ^22^. It is also the world’s largest multimodal imaging study, including brain imaging data ^25^. In the present study, we used resting-state functional magnetic resonance imaging (rfMRI) data from UK Biobank to assess resting-state functional connectivity within a connectome relevant to interoception. UK Biobank’s imaging protocols can be accessed online (http://biobank.ctsu.ox.ac.uk/crystal/refer.cgi?id=2367). Further information has been published elsewhere regarding brain imaging in UK Biobank ^26^ and the processing pipeline ^27^. rfMRI data was acquired at three identical Biobank imaging centres across the UK, using a 3T Siemens Skyra scanner with a 32-channel head coil. The T2*-weighted images with BOLD contrast were measured with gradient-echo echo-planar imaging (GE-EPI) with x8 multi-slice (multiband) acceleration, no in-plane acceleration, flip angle = 52°, fat saturation, TR = 735ms, TE = 39ms, FOV = 88 × 88 × 64, voxel size = 2.4mm (isotropic). Resting-state scans lasted 6 minutes, comprising 490 time points.

#### rfMRI processing

We analysed data that had undergone standard pre-processing, as follows: As described in the imaging documentation, rfMRI data were pre-processed using FSL (the FMRIB Software Library, https://fsl.fmrib.ox.ac.uk/fsl/fslwiki). Motion correction MCFLIRT ^28^, grand-mean intensity normalisation, high-pass temporal filtering (Gaussian-weighted least-squares straight line fitting, sigma = 50s), EPI unwarping and gradient distortion correction (GDC) were applied. EPI and GDC unwarping both included alignment to the T1 structural image using FLIRT ^28,29^. Images were nonlinearly warped to MNI152 standard space utilising FNIRT ^30,31^. Finally, structured artefacts were removed using independent-component analysis (ICA) followed by FMRIB’s ICA-based X-noiseifier ^32–34^, resulting in the FIX-cleaned timeseries, which were used to compute the functional connectivity matrix within our network of interest.

#### Imaging-derived phenotypes (IDPs)

We created binary region of interest masks (see Table 1 for additional information) for eleven regions representing the nodes of a network relevant to interoception. The nodes included the insular cortex (IC) ^7,17,35–37^, functionally differentiated into a posterior (pIC) and anterior (aIC) section. The latter is often co-activated with anterior cingulate cortex (ACC) ^17,38^, which can functionally be subdivided into a ventral (vACC) ^39–41^ and dorsal (dACC) portion. dACC can functionally be associated with IC and sensorimotor structures ^35,42^, where we selected primary sensory (S1) and motor (M1) cortex. Nodes further included ventromedial prefrontal cortex (vmPFC) situated next to vACC ^43^ and dorsomedial prefrontal cortex (dmPFC), which can be co-activated with adjacent dACC and IC ^42^. Finally, we included subcortical amygdala (Amy) ^44–46^ which shows extensive reciprocal connections with IC ^47^, and the periaqueductal grey (PAG), where lateral (lPAG) and ventrolateral (vlPAG) subregions have been differentiated ^48,49^.

**Table 1.**
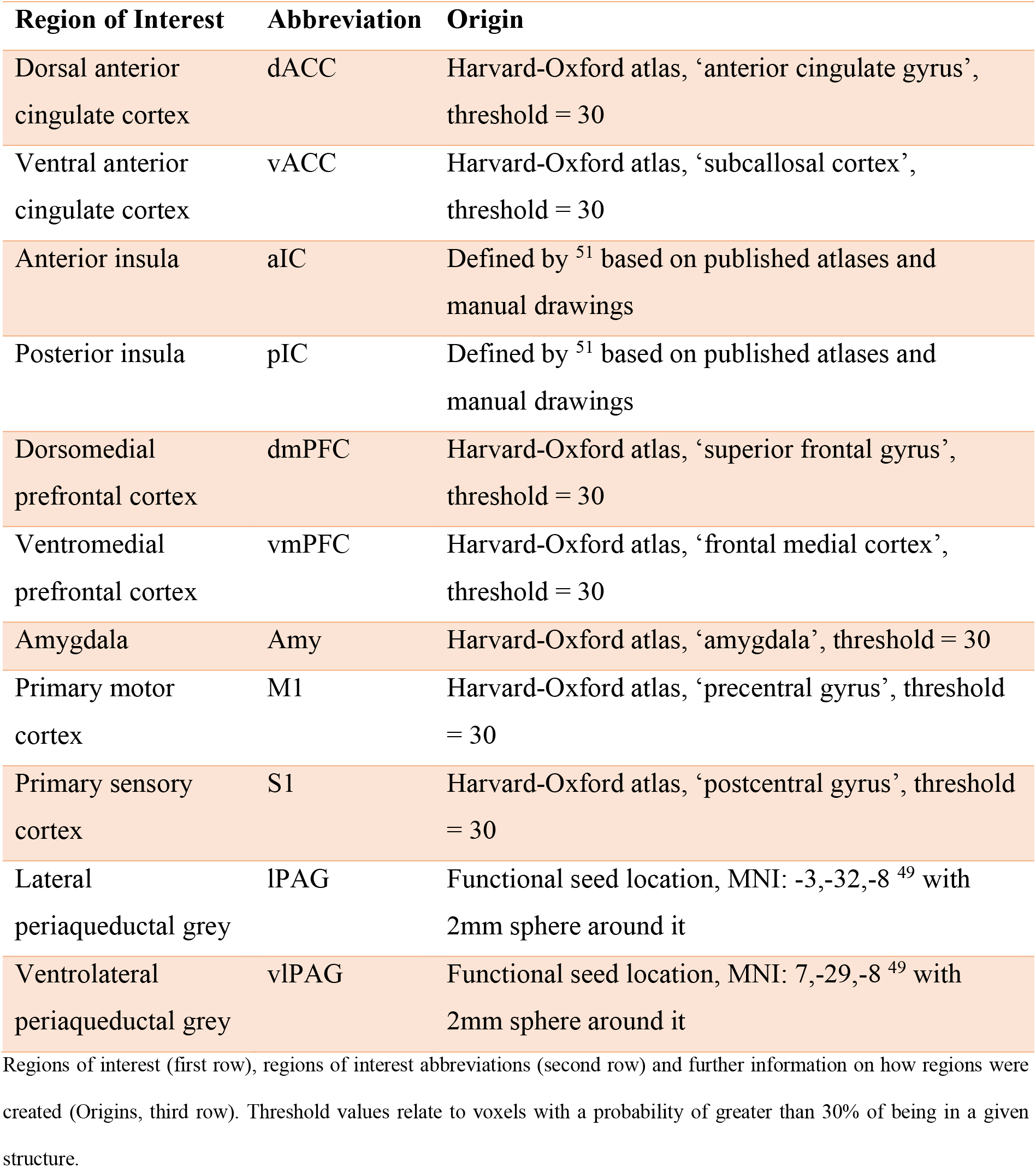

The 4-D timeseries were extracted within each region of interest mask by averaging over voxels. Functional connectivity measures were obtained by performing partial correlations between the 11 region of interest specific timeseries, to generate 55 imaging-derived phenotypes (IDPs), which correspond to one half of the correlation matrix resulting from 11 regions of interest. IDPs were then winsorised at 1^st^ and 99^th^ percentiles to smooth over outliers, standardised and deconfounded by regressing effects of age, sex, imaging centre, head motion, head size and table position out of the winsorised, normalised IDPs. By winsorising the data, extreme values do not have to be removed from the dataset, but instead are trimmed. In this case, 1% of the extreme values (positive and negative) were set to the values of the 1^st^ and 99^th^ percentile ^50^. Standardisation of the data meant that for each variable, the mean was 0 and the standard deviation was 1.

### Non-imaging derived phenotypes

Non-imaging derived phenotypes (nIDPs) were a set of 170 UK Biobank measures (see supplemental information table 1 for more details, and supplemental information table 2 for excluded variables) that broadly covered physiological/physical health (including also breathing, body measures and bloods), mental health and well-being (also including cognition) and lifestyle (including physical activity, smoking, nutrition, and job). Data were winsorised at 1^st^ and 99^th^ percentiles. Missing data were imputed (no variables had more than 50% of missing data, median: 1.98%, interquartile range: 3.62%) using k-nearest (k=1) neighbour imputation. nIDPs were normalised and deconfounded (same as IDPs).

### Planned statistical analyses

First, principal component analysis was performed separately on IDPs (55 brain connectivity variables) and nIDPs (168 variables), identifying the top components that cumulatively explained 80% of the variance. These principal components were submitted to canonical correlation analysis (CCA) using MATLAB’s *canoncorr* function. CCA is a multivariate technique to detect modes of covariation between two sets of variables, in this case the two sets were IDPs and nIDPs ^26,52^. Modes represent linear combinations of the variables in each set that maximally covary across subjects, termed *canonical variates*. CCA computes as many modes as there are variables in the smaller set. To identify statistically significant modes (alpha level < 0.05 with family-wise correction for multiple comparisons), a null distribution was established with 100,000 permutations of the subject rows of one variable set relative to the other variable set. The strongest correlation from each permutation was selected to establish the null distribution. Modes of covariation can be interpreted by inspecting the weight of each given variable as determined by the correlation coefficients between the individual variables in a set and their canonical variate. We took a conservative approach to the interpretation of these four modes by creating averages of the canonical variates *U* and *V* and correlating these averaged variates with the (non PCA-reduced) values for the IDPs and nIDPs. To facilitate interpretation, only correlation coefficients ≥ 0.2 and ≤ -0.2 were then reported ^52^.

## Results

After PCA, the top 33 components related to IDPs cumulatively explained 81.38% of the variance. The top 88 components related to nIDPs explained 80.37% of the variance. Those scores were used to calculate the canonical variates and resulting modes of covariation. Four modes of covariation were found to be statistically significant after permutation testing (Mode 1 canonical correlation: 0.2351, p<0.0001, r^2^=0.0553; Mode 2 canonical correlation: 0.1512, p<0.0001, r^2^=0.0229; Mode 3 canonical correlation: 0.1336, p<0.0001, r^2^=0.0178; Mode 4 canonical correlation: 0.1210, p<0.0001, r^2^=0.0146; figure 1).

**Figure 1:**
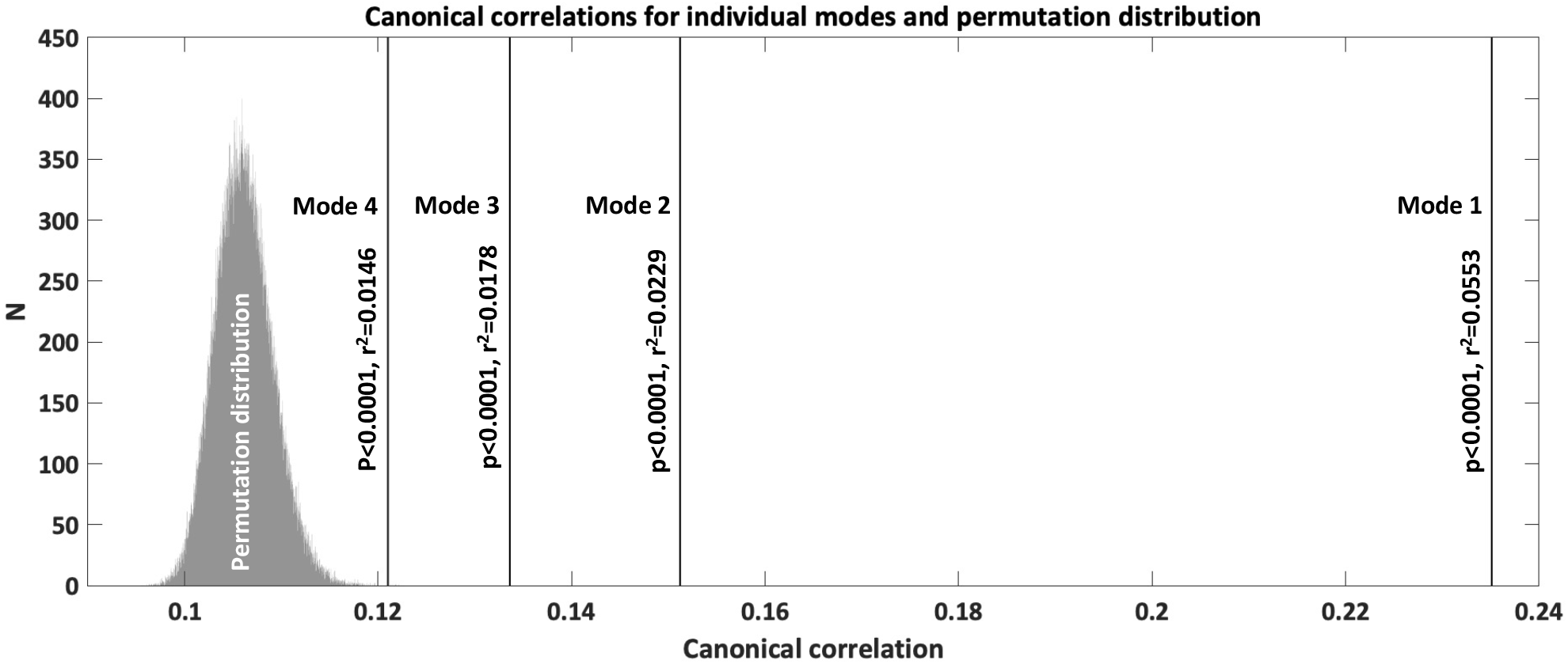
Canonical correlations for the four significant modes of co-variation and permutation distribution. Canonical correlations for individual modes were evaluated against a null distribution of canonical correlations (grey bars) derived from permuting subject rows of one variable set relative to the other variable set 100 000 times. The strongest correlation from each permutation was selected to establish the permutation distribution. Vertical lines mark canonical correlations for the four modes of the original data which were significantly different from the null distribution (statistical significance and variance explained are shown).

We labelled Mode 1 (figure 2: Mode 1) explaining 5.53% of the variance as related to affect and cardiovascular health, since nIDPs may primarily be associated with affective processing including also cardiovascular variables. The neural network of mode 1 was very dense and characterised by prominent involvement of amygdala (3 connections), IC (4 connections), dACC (5 connections) and somatosensory and motor cortices (S1/M1, 8 connections).

**Figure 2:**
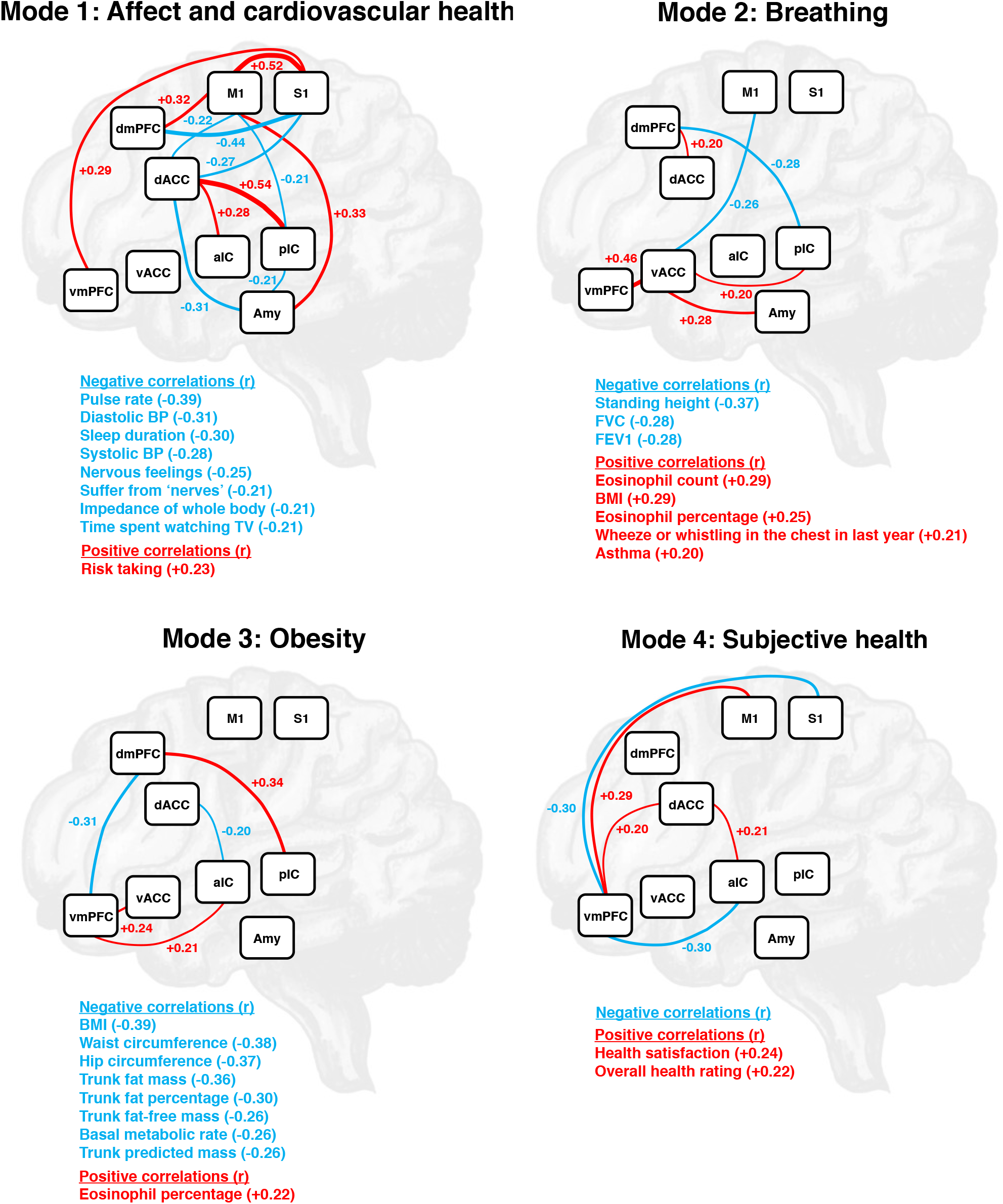
Significant modes: Neural circuits and associated non-imaging related phenotypes. Mode 1: Affect and cardiovascular health, p < 0.0001, variance explained: 5.53%. Mode 2: Breathing, p < 0.0001, variance explained: 2.29%. Mode 3: Obesity, p < 0.0001, variance explained: 1.78%. Mode 4: Subjective health, p < 0.0001, variance explained: 1.46%. Negative correlations with canonical variates are depicted in blue, positive correlations in red. Abbreviations: BMI (body mass index), BP (blood pressure), FEV1 (forced expiratory volume in 1s), FVC (forced vital capacity), vmPFC (ventromedial prefrontal cortex), dmPFC (dorsomedial prefrontal cortex), vACC (ventral anterior cingulate cortex), dACC (dorsal anterior cingulate cortex), aIC (anterior insular cortex), pIC (posterior insular cortex), Amy (amygdala), M1 (primary motor cortex), S1 (primary sensory cortex).

Mode 2 (figure 2: Mode 2) explaining 2.29% of the variance was labelled as related to breathing, since the set of nIDPs may predominantly be linked to breathing and respiratory disease. The neural network of mode 2 was centred at vACC (4 connections) with a comparably strong vACC-vmPFC connection. pIC and dmPFC had each 2 connections.

We labelled Mode 3 (figure 2: Mode 3) explaining 1.78% of the variance as being associated mainly with obesity (and metabolism). The neural network underlying mode 3 included all prefrontal, insular and anterior cingulate cortices. dmPFC featured 2 and vmPFC 3 connections.

Mode 4 (figure 2: Mode 4) which explained 1.46% of the variance contained nIDPs of subjective health and was labelled accordingly. The neural network of mode 4 contained predominantly vmPFC connections to somatosensory, motor, dorsal anterior cingulate and anterior insular cortices. dACC featured 2 connections.

## Discussion

In 19,020 participants we identified four statistically independent modes of co-variation between connectivity profiles within interoceptive brain regions and measures of lifestyle and health related functioning. Modes broadly represented distinct functional domains: Mode 1 related to affect and cardiovascular health, Mode 2 to breathing, Mode 3 to obesity and Mode 4 to subjective health. Modes go far beyond simple region-function associations and represent specific “neural fingerprints” of individual connectivity profiles associated with functional domains in the context of interoception. Our research contributes to the conceptualisation of interoception and sets the scope for future more directed and experimental research on the neural correlates of interoceptive involvement in various lifestyle and health related variables.

There is a large correspondence between resting-state and task-related networks ^53^, and resting state activity can shape task activations ^54^ and predict behaviour ^55^. By not employing a specific task, we could relate functional connectivity to different health-related fields in one integrative analysis, delineating shared and separated mechanisms. Consequently, our research may contribute to a biologically informed, brain-based characterisation of co-morbid conditions.

For instance, Mode 1 shows a brain network linking variables of negative affect with variables of cardiovascular health, supporting previous research suggesting a link between these dimensions at the neural level ^56^. Furthermore, breathing pathology has been related to obesity ^57,58^ and negative affect ^57,59,60^. Dimensions of affect, breathing and obesity were to a large extent dissociated into mutually uncorrelated modes and different phenotypes of pathology may differ in the involvement of individual mode dimensions. However, breathing Mode 2 still contained variables arguably associated with obesity (BMI) and potentially affective processing (vACC-Amygdala).

Affective processing related to anxiety and depression ^61,62^ (Mode 1), breathing ^60^ (Mode 2), obesity and metabolic function ^16^ (Mode 3), and subjective health (Mode 4) and well-being ^20,63,64^ have previously been linked to interoception, and we suggest that further studies should consider their relation to interoception with regard to the different neural circuits highlighted by the present research.

As circuits share contributions of critical interoceptive and domain general insular, anterior cingulate and medial prefrontal regions, pathological processing in these regions may have an impact on more than one functional domain. This could better explain why pathologies occur as co-morbidities, and why interoception is relevant for a large variety of disorders ^20,65^. Our study could thus give rise to future research on a neurally informed classification of co-morbid conditions and subjective health in the context of interoception cutting across traditional diagnostic boundaries ^66^.

Neural circuits of individual modes were perfectly dissociated between modes 1 and 2, and were generally specific representing distinct neural fingerprints of functional domains. This contributes to research on the conceptualisation of interoception, which describes and seeks to clarify relationships between different interoceptive functions associated with more lower level sensing and regulation of bodily states and higher level emotional, cognitive or even self-related processing ^9,16^. In the following, we will consider each mode individually.

### Mode 1 – Affect and cardiovascular health

Mode 1 explained 5.53% of the variance and may primarily be associated with affective processing and cardiovascular health. Negative correlations with canonical variates were seen for the following nIDPs related to anxiety and depression, i.e. negative affect ^62^: Unfavourable nervous perceptions (‘Nervous feelings’ / ‘Suffer from nerves’), ‘Sleep duration’ ^67,68^, ‘Pulse rate’ ^69^ and ‘Time spent watching TV’ ^70,71^. ‘Diastolic BP’ and ‘Systolic BP’ (diastolic and systolic blood pressure) correlated also negatively with canonical variates, where positive and negative relationships with negative affect have been reported ^72–74^. We obtained a plausible inverse relationship of variables of negative affect with ‘Risk taking’, as risk aversion has been related to negative affect ^75,76^.

We found a network including dACC, dmPFC, IC and sensorimotor connections which were much more extensive than in any other mode. This seems plausible, as dACC, dmPFC, IC and sensorimotor cortices are specifically important for the generation of interoceptive awareness ^35,42,77^, and the perception of interoceptive input has been theorised to be crucial for affective experience ^5–10^. Moreover, amygdala connections were mainly found in this mode. Amygdala is critical for emotional processing, especially the processing of fear ^45,46^, in particular within a network containing amygdala, IC and/or ACC ^78–81^, which may form a circuit of negative affect ^82^ and were linked to variables of negative affect in mode 1.

Although our findings cannot clarify causal relationships, they are well in line with other research linking both negative emotions and variables of cardiovascular health to mPFC, ACC, IC and amygdala (see Kraynak et al. ^56^ for a review). We provide explicit connections between these regions at the population level that should inform future investigations examining the visceromotor and viscerosensory mechanisms linking affective states and physiological changes.

### Mode 2 - Breathing

We labelled Mode 2 explaining 2.29% of the variance as related to breathing. Major variables of respiratory health ‘FVC’ (forced vital capacity) and ‘FEV1’(forced expiratory volume in 1s) ^83^ were negatively associated with canonical variates. ‘Standing height’ showed the same relationship, which is plausible given that taller subjects can exhale larger volumes. Variables related to respiratory disease included ‘Wheeze or whistling in the chest in the last year’, ‘Asthma’, ‘Eosinophil count’, ‘Eosinophil percentage’ and ‘BMI’ (Body mass index) and showed a meaningful positive correlation with canonical variates. Eosinophils promote inflammation and are increased in some types of asthma ^84^. Obesity has been shown to be a major risk factor and disease modifier in asthma, where obese asthmatics show more frequent and severe exacerbations ^58^.

vACC connections were particularly prominent in Mode 2. A vACC-vmPFC connection was especially strong and linked to variables of breathing pathology. This supports previous research in the context of chronic obstructive pulmonary disease, where vmPFC and vACC were associated with the evaluation of breathlessness and suggested to contribute to a poor correlation between lung function and symptoms ^85^. vACC-amygdala connectivity was also linked to breathing pathology in Mode 2 and may be a target for future research, as amygdala processing has been associated with breathing inhibition ^86^, and functional connections including amygdala and vACC are implicated in pathological emotional processing ^87,88^.

### Mode 3 – Obesity

Mode 3 explaining 1.78% of the variance was mainly related to obesity (and metabolism). ‘BMI’, ‘Waist circumference’, ‘Hip circumference’, ‘Trunk fat mass’, ‘Trunk fat percentage’, and ‘Basal metabolic rate’ were all part of mode 3, correlating negatively with canonical variates. ‘Trunk predicted mass’ and ‘Trunk fat-free mass’ correlated also negatively with canonical variates, where an association of body fat and fat-free mass has been reported previously ^89^. Eosinophils (‘Eosinophil percentage’) which are implicated in tissue homeostasis ^90^ correlated positively with canonical variates, but their relationship with obesity is under debate ^91,92^.

Physiological functioning requires the sensing, interpretation and control of energy-status-related internal states, integrating them with energy needs, learned experiences and exteroceptive information which motivates behavioural responses such as feeding behaviour ^16,93^. Therefore, it seems plausible that mode 3 was characterised by a network including pIC, the major substrate of viscerosensation ^17,36,37^, vACC and aIC, the primary regions of visceromotor control ^17,65^, and dorsal and ventral medial prefrontal cortices controlling affective and cognitive processes and behaviour ^43,94,95^.

Functional connectivity modulations in brain structures including vmPFC, dmPFC, dACC and aIC were shown in obese patients following bariatric surgery ^96^, and our research suggests that a network including these regions may play a more general role in obesity.

A negative relationship of BMI and resting-state connectivity has been shown for several regions in obese patients ^97^, and we confirm this relationship for dmPFC-pIC connectivity, which represented the strongest connection of Mode 3.

### Mode 4 – Subjective Health

Mode 4 explained 1.46% of the variance and was related to variables of subjective health including ‘Health satisfaction’ and ‘Overall health rating’. Variables of subjective health should involve processes related to self-reference, appraisal and emotion. Crucially, vmPFC was the centre of the network of mode 4 and connected to every other region of the network. This seems plausible, as many different functions that have been associated with vmPFC have been summarised to relate to subjective value estimation ^98^, specifically in relation to the self ^99^, and the contextual shaping of affective information ^43^. Subjective health is an important indicator to describe the actual needs and problems of patients and can reflect and influence physiology. Interestingly, vmPFC plays a central role in mediating the interplay between self-related, also interoceptive states and physiology ^100^, and we provide explicit vmPFC connections for future more directed research.

### Strengths and limitations

The major strength of the study is the sample size of more than 19,000 participants ^25^ which has allowed us to obtain meaningful and robust hidden associations between two sets of variables using CCA ^101,102^. Our data-driven approach enabled a truly exploratory analysis linking specific brain networks with a possible relation to interoception to a broad range of specific correlated lifestyle and health related variables. Although our sample size facilitates the detection of small effects, the four modes identified together represent more than 10% of population variance. The identification of multiple significant modes of population variance was not a foregone conclusion. For example, Smith et al. ^23^ who used CCA to analyse multivariable whole brain connectivity and diverse health relevant non-brain data in 461 subjects from the Human Connectome Project obtained only one strong mode.

The present large-scale dataset is observational and cross-sectional, so that causal interpretations of our results are not possible. In addition, the dataset does not contain explicit measures of interoception. As brain regions relevant to interoception are also relevant to other functions, we cannot draw definite conclusions about interoception ^103^. However, we relate specific connectivity profiles within interoceptive brain regions to specific health related domains. These domain specific neural fingerprints can contribute to conceptualizing interoception ^9,16^ and inform future experimental interoceptive research.

We included lPAG and vlPAG as previous research suggests that the PAG is a brain structure involved in interoception, information integration and autonomic behavioural control, crucially in the context of breathing ^48,49^. However, we did not show resting PAG connectivity with breathing mode 2 or any other mode. As lPAG and vlPAG ROIs were necessarily very small, a low signal-to-noise ratio may have prevented the detection of an effect.

Our results are inevitably constrained by the selection of variables. We pre-selected a set of areas identified in previous studies relevant to interoception. Non-imaging phenotypes were selected very broadly to chart neural variables of various lifestyle and health related conditions. Due to the breadth of variables and novelty of our approach variable selection was necessarily subject to educated judgement. Upcoming investigations should therefore aim to replicate and extend our findings employing variations in the selection of imaging and non-imaging related variables.

## Conclusion

To conclude, we studied the relationship of a functional network of brain regions relevant to interoception to a broad selection of non-imaging health and lifestyle related phenotypes in more than 19.000 UK Biobank study participants. Our integrative and data driven analysis revealed four modes of population co-variation with distinct neural circuits which could respectively meaningfully be associated with affect and cardiovascular health, breathing, obesity or subjective health and may also relate to interoception. Circuits can be regarded as specific neural “fingerprints” of functional domains and set the scope for more directed research on the neurobiology of interoceptive involvement in different lifestyle and health related factors. Therefore, our research contributes to the conceptualisation of interoception and may help to better understand co-morbid conditions in the light of shared interoceptive structures.

## Acknowledgements

We want to thank UK Biobank for providing the data analysed by the present research (data access application number: 25673).

