## Supplemental tables 1 and 2 for "Functional connectivity between interoceptive brain regions is associated with distinct health-related domains - a population-based neuroimaging study"

### Supplemental information

#### Supplemental table 1: Full list of 170 selected nIDPs

| Category | Variable name | Biobank code |
|---|---|---|
| **Physiological health** | | |
| *Various* | Pulse rate, automated reading | 102 |
| | Number of self-reported cancers | 134 |
| | Number of self-reported non-cancer illnesses | 135 |
| | Number of operations, self-reported | 136 |
| | Long-standing illness, disability or infirmity | 2188 |
| | Chest pain or discomfort | 2335 |
| | Diabetes diagnosed by doctor | 2443 |
| | Cancer diagnosed by doctor | 2453 |
| | Fractured/broken bones in last 5 years | 2463 |
| | Other serious medical condition/disability diagnosed by doctor | 2473 |
| | Diastolic blood pressure, automated reading | 4079 |
| | Systolic blood pressure, automated reading | 4080 |
| *Breathing* | Wheeze or whistling in the chest in last year | 2316 |
| | Peak expiratory flow (PEF) | 3064 |
| | Shortness of breath walking on level ground | 4717 |
| | asthma | 6152 |
| | blood clot in leg | 6152 |
| | blood clot in lung | 6152 |
| | Hayfever, allergic rhinitis or eczema | 6152 |
| | Forced expiratory volume in 1-second (FEV1), Best measure | 20150 |
| | Forced vital capacity (FVC), Best measure | 20151 |
| | Cough on most days | 22502 |
| | Years of cough on most days | 22503 |
| | Bring up phlegm/sputum/mucus on most days | 22504 |
| | Years of bringing up phlegm/sputum/mucus on most days | 22505 |
| | FEV1/FVC ratio | calculated |
| *Body measures* | Waist circumference | 48 |
| | Hip circumference | 49 |
| | Standing height | 50 |
| | Body mass index (BMI) | 21001 |
| | Basal metabolic rate | 23105 |
| | Impedance of whole body | 23106 |
| | Trunk fat percentage | 23127 |
| | Trunk fat mass | 23128 |

| | | |
|---|---|---|
| | Trunk fat-free mass | 23129 |
| | Trunk predicted mass | 23130 |
| *Bloods* | White blood cell (leukocyte) count | 30000 |
| | Red blood cell (erythrocyte) count | 30010 |
| | Haemoglobin concentration | 30020 |
| | Haematocrit percentage | 30030 |
| | Mean corpuscular volume | 30040 |
| | Mean corpuscular haemoglobin | 30050 |
| | Mean corpuscular haemoglobin concentration | 30060 |
| | Red blood cell (erythrocyte) distribution width | 30070 |
| | Platelet count | 30080 |
| | Platelet crit | 30090 |
| | Mean platelet (thrombocyte) volume | 30100 |
| | Platelet distribution width | 30110 |
| | Lymphocyte count | 30120 |
| | Monocyte count | 30130 |
| | Neutrophill count | 30140 |
| | Eosinophill count | 30150 |
| | Basophill count | 30160 |
| | Nucleated red blood cell count | 30170 |
| | Lymphocyte percentage | 30180 |
| | Monocyte percentage | 30190 |
| | Neutrophill percentage | 30200 |
| | Eosinophill percentage | 30210 |
| | Basophill percentage | 30220 |
| | Nucleated red blood cell percentage | 30230 |
| | Reticulocyte percentage | 30240 |
| | Reticulocyte count | 30250 |
| | Mean reticulocyte volume | 30260 |
| | High light scatter reticulocyte percentage | 30290 |
| | High light scatter reticulocyte count | 30300 |
| **Mental health and well-being** | | |
| *Various* | Mood swings | 1920 |
| | Miserableness | 1930 |
| | Irritability | 1940 |
| | Sensitivity / hurt feelings | 1950 |
| | Fed-up feelings | 1960 |
| | Nervous feelings | 1970 |
| | Worrier / anxious feelings | 1980 |
| | Tense / ''highly strung'' | 1990 |
| | Worry too long after embarrassment | 2000 |
| | Suffer from ''nerves'' | 2010 |

| | | |
|---|---|---|
| | Loneliness, isolation | 2020 |
| | Guilty feelings | 2030 |
| | Risk taking | 2040 |
| | Frequency of depressed mood in last 2 weeks | 2050 |
| | Frequency of unenthusiasm / disinterest in last 2 weeks | 2060 |
| | Frequency of tenseness / restlessness in last 2 weeks | 2070 |
| | Frequency of tiredness / lethargy in last 2 weeks | 2080 |
| | Seen doctor (GP) for nerves, anxiety, tension or depression | 2090 |
| | Seen a psychiatrist for nerves, anxiety, tension or depression | 2100 |
| | Able to confide | 2110 |
| | Overall health rating | 2178 |
| | Happiness | 4526 |
| | Work/job satisfaction | 4537 |
| | Health satisfaction | 4548 |
| | Family relationship satisfaction | 4559 |
| | Friendships satisfaction | 4570 |
| | Financial situation satisfaction | 4581 |
| | Ever depressed for a whole week | 4598 |
| | Longest period of depression | 4609 |
| | Number of depression episodes | 4620 |
| | Ever unenthusiastic/disinterested for a whole week | 4631 |
| | Ever manic/hyper for 2 days | 4642 |
| | Ever highly irritable/argumentative for 2 days | 4653 |
| | Neuroticism score | 20127 |
| *Cognition* | Number of correct matches in round | 398 |
| | Number of incorrect matches in round | 399 |
| | Time to complete round | 400 |
| | Time to answer | 4288 |
| | Number of attempts | 4291 |
| | Fluid intelligence score | 20016 |
| | Prospective memory result | 20018 |
| | Mean time to correctly identify matches | 20023 |
| | Fluid intelligence questions attempted within time limit | 20128 |
| **Lifestyle** | | |
| *Physical activity* | Number of days/week walked 10+ minutes | 864 |
| | Duration of walks | 874 |
| | Number of days/week of moderate physical activity 10+ minutes | 884 |
| | Duration of moderate activity | 894 |
| | Number of days/week of vigorous physical activity 10+ minutes | 904 |
| | Duration of vigorous activity | 914 |
| | Usual walking pace | 924 |
| | Frequency of stair climbing in last 4 weeks | 943 |

| | | |
|---|---|---|
| | Frequency of walking for pleasure in last 4 weeks | 971 |
| | Duration walking for pleasure | 981 |
| | Frequency of light DIY in last 4 weeks | 1011 |
| | Duration of light DIY | 1021 |
| | Time spent outdoors in summer | 1050 |
| | Time spent outdoors in winter | 1060 |
| | Time spent watching television | 1070 |
| | Time spent using computer | 1080 |
| | Time spent driving | 1090 |
| | Drive faster than motorway speed limit | 1100 |
| | Sleep duration | 1160 |
| | Frequency of heavy DIY in last 4 weeks | 2624 |
| | Duration of heavy DIY | 2634 |
| | Frequency of other exercises in last 4 weeks | 3637 |
| | Duration of other exercises | 3647 |
| *Smoking* | Current tobacco smoking | 1239 |
| | Past tobacco smoking | 1249 |
| | Smoking/smokers in household | 1259 |
| | Exposure to tobacco smoke at home | 1269 |
| | Exposure to tobacco smoke outside home | 1279 |
| | Smoking status | 20116 |
| | Ever smoked | 20160 |
| *Nutrition* | Cooked vegetable intake | 1289 |
| | Salad / raw vegetable intake | 1299 |
| | Fresh fruit intake | 1309 |
| | Dried fruit intake | 1319 |
| | Oily fish intake | 1329 |
| | Non-oily fish intake | 1339 |
| | Processed meat intake | 1349 |
| | Poultry intake | 1359 |
| | Beef intake | 1369 |
| | Lamb/mutton intake | 1379 |
| | Pork intake | 1389 |
| | Cheese intake | 1408 |
| | Bread intake | 1438 |
| | Cereal intake | 1458 |
| | Tea intake | 1488 |
| | Coffee intake | 1498 |
| | Water intake | 1528 |
| | Alcohol intake frequency | 1558 |
| | Average weekly red wine intake | 1568 |
| | Average weekly champagne plus white wine intake | 1578 |
| | Average weekly beer plus cider intake | 1588 |

| | | |
|---|---|---|
| | Average weekly spirits intake | 1598 |
| | Average weekly fortified wine intake | 1608 |
| | Alcohol usually taken with meals | 1618 |
| | Alcohol drinker status | 20117 |
| *Job* | Townsend deprivation index at recruitment | 189 |
| | Time employed in main current job | 757 |
| | Length of working week for main job | 767 |
| | Job involves mainly walking or standing | 806 |
| | Job involves heavy manual or physical work | 816 |
| | Job involves shift work | 826 |
| | Age completed full time education | 845 |

If variables featured more than one data point the most recent data point was selected or, if appropriate, an average measure was calculated. In some cases, variables were re-coded for consistency and to fit the analysis structure. Unclear information from categories of individual variables (e.g. “Prefer not to say”, “Do not know”) was excluded from the analysis.

**Supplemental table 2: Full list of 107 excluded nIDPs**

| **Reason for exclusion** | **Variable name** | **Biobank code** |
|---|---|---|
| **Used for unconfounding** | Year of birth | 34 |
| | Month of birth | 52 |
| | Scan date | 53 |
| | Sex | 31 |
| **>50% missing data** | Frequency of strenuous sports in last 4 weeks | 991 |
| | Duration of strenuous sports | 1001 |
| | Light smokers, at least 100 smokes in lifetime | 2644 |
| | Age started smoking in former smokers | 2867 |
| | Number of cigarettes previously smoked daily | 2887 |
| | Age stopped smoking | 2897 |
| | Ever stopped smoking for 6+ months | 2907 |
| | General pain for 3+ months | 2956 |
| | Neck/shoulder pain for 3+ months' | 3404 |
| | Hip pain for 3+ months | 3414 |
| | Job involve night shift work | 3426 |

| | | |
|---|---|---|
| | Age started smoking in current smokers | 3436 |
| | Number of cigarettes currently smoked daily (current cigarette smokers) | 3456 |
| | Back pain for 3+ months | 3571 |
| | Chest pain or discomfort walking normally | 3606 |
| | Chest pain due to walking ceases when standing still | 3616 |
| | Chest pain or discomfort when walking uphill or hurrying | 3751 |
| | Knee pain for 3+ months | 3773 |
| | Age asthma diagnosed | 3786 |
| | Headaches for 3+ months | 3799 |
| | Facial pains for 3+ months | 4067 |
| | Maximum digits remembered correctly | 4282 |
| | Longest period of unenthusiasm / disinterest | 5375 |
| | Number of unenthusiastic/disinterested episodes | 5386 |
| | Length of longest manic/irritable episode | 5663 |
| | Severity of manic/irritable episodes | 5674 |
| | Doctor restricts physical activity due to heart condition | 6014 |
| | Chest pain felt during physical activity | 6015 |
| | Chest pain felt outside physical activity | 6016 |
| | Able to walk or cycle unaided for 10 minutes | 6017 |
| | Maximum workload during fitness test | 6032 |
| | Maximum heart rate during fitness test | 6033 |
| | Single episode of probable major depression | 20123 |
| | Probable recurrent major depression (moderate) | 20124 |
| | Probable recurrent major depression (severe) | 20125 |
| | Bipolar and major depression status | 20126 |
| | Age of stopping smoking | 22507 |
| | Amount of tobacco currently smoked | 22508 |
| **<1% confirmed events** | Used an inhaler for chest within last hour | 3090 |
| | Average weekly intake of other alcoholic drinks | 5364 |
| | Previously smoked cigarettes on most/all days | 5959 |
| | Emphysema/chronic bronchitis | 6152 |
| | Number of cigarettes previously smoked daily (current cigar/pipe smoker) | 6183 |
| | Age stopped smoking cigarettes (current cigar/pipe or previous cigarette smoker) | 6194 |
| | Doctor diagnosed emphysema | 22128 |
| | Doctor diagnosed chronic bronchitis | 22129 |
| | Doctor diagnosed COPD (chronic obstructive pulmonary disease) | 22130 |
| | Doctor diagnosed cystic fibrosis | 22131 |
| | Doctor diagnosed alpha-1 antitrypsin deficiency | 22132 |
| | Doctor diagnosed sarcoidosis | 22133 |
| | Doctor diagnosed bronchiectasis | 22134 |

| | | |
|---|---|---|
| | Doctor diagnosed idiopathic pulmonary fibrosis | 22135 |
| | Doctor diagnosed fibrosing alveolitis/unspecified alveolitis | 22136 |
| | Doctor diagnosed tuberculosis | 22137 |
| | Doctor diagnosed silicosis | 22138 |
| | Doctor diagnosed asbestosis | 22139 |
| | Doctor diagnosed lung cancer (not mesothelioma) | 22140 |
| | Doctor diagnosed mesothelioma of the lung | 22141 |
| **Duplicates** | Systolic blood pressure, manual reading | 93 |
| | Diastolic blood pressure, manual reading | 94 |
| | Pulse rate (during blood-pressure measurement) | 95 |
| | Forced vital capacity (FVC) | 3062 |
| | Forced expiratory volume in 1-second (FEV1) | 3063 |
| | Average monthly red wine intake | 4407 |
| | Average monthly champagne plus white wine intake | 4418 |
| | Average monthly beer plus cider intake | 4429 |
| | Average monthly spirits intake | 4440 |
| | Average monthly fortified wine intake | 4451 |
| | Average monthly intake of other alcoholic drinks | 4462 |
| | Doctor diagnosed hayfever or allergic rhinitis | 22126 |
| | Doctor diagnosed asthma | 22127 |
| | Year of birth | 22200 |
| | Tobacco smoking | 22506 |
| | BMI | 23104 |
| **Postcode dependent variables *** | Frequency of travelling from home to job workplace | 777 |
| | Distance between home and job workplace | 796 |
| | Frequency of friend/family visits | 1031 |
| | Nitrogen dioxide air pollution; 2010 | 24003 |
| | Nitrogen oxides air pollution; 2010 | 24004 |
| | Particulate matter air pollution (pm10); 2010 | 24005 |
| | Particulate matter air pollution (pm2.5); 2010 | 24006 |
| | Particulate matter air pollution (pm2.5) absorbance; 2010 | 24007 |
| | Particulate matter air pollution 2.5-10um; 2010 | 24008 |
| | Traffic intensity on the nearest road | 24009 |
| | Inverse distance to the nearest road | 24010 |
| | Traffic intensity on the nearest major road | 24011 |
| | Inverse distance to the nearest major road | 24012 |
| | Total traffic load on major roads | 24013 |
| | Close to major road | 24014 |
| | Sum of road length of major roads within 100m | 24015 |
| | Nitrogen dioxide air pollution; 2005 | 24016 |
| | Nitrogen dioxide air pollution; 2006 | 24017 |
| | Nitrogen dioxide air pollution; 2007 | 24018 |

| | | |
|---|---|---|
| | Particulate matter air pollution (pm10); 2007 | 24019 |
| **Structural brain variables **** | Volumetric scaling from T1 head image to standard space | 25000 |
| | Volume of peripheral cortical grey matter (normalised for head size) | 25001 |
| | Volume of peripheral cortical grey matter | 25002 |
| | Volume of ventricular cerebrospinal fluid (normalised for head size) | 25003 |
| | Volume of ventricular cerebrospinal fluid | 25004 |
| | Volume of grey matter (normalised for head size) | 25005 |
| | Volume of grey matter | 25006 |
| | Volume of white matter (normalised for head size) | 25007 |
| | Volume of white matter | 25008 |

* Postcode dependent variables may not be representative as moving houses can strongly influence these variables.
** Structural brain variables were not the focus of the present investigation.
